# Deletion of *TUBB4A* mitigates the oligodendrocyte and neuronal deficits in human iPSCs derived from individuals affected by H-ABC

**DOI:** 10.64898/2026.07.31.742062

**Authors:** Zhongqi Alice Hou, Rolf Dale Bates, Prabhat R Napit, Luis Garcia, Anjali Bhagavatula, Julia L Hacker, Teshawn Johnson, Alyssa Gagne, Jean Ann Maguire, Asako Takanohashi, Deborah French, Akshata Almad, Judith Grinspan, Adeline Vanderver, Sunetra Sase

## Abstract

*TUBB4A*-related leukodystrophy (*TUBB4A*-LD) is a rare neurologic disorder with a broad spectrum of phenotypes, including severe early infantile encephalopathy, late infantile Hypomyelination with Atrophy of the Basal ganglia and Cerebellum (H-ABC), and milder late infantile forms. H-ABC is closely associated with a recurrent pathogenic variant, p.Asp249Asn, in the gene encoding tubulin beta class IVA (TUBB4A), a microtubule component. H-ABC presents with progressive dystonia, mobility loss, aphasia, and swallowing dysfunction in childhood. H-ABC results in cell-autonomous deficits in oligodendrocytes (OLs), cerebellar granule neurons, and medium spiny neurons (MSNs). Antisense oligonucleotides targeting *Tubb4a* can alleviate symptoms in H-ABC mouse models. However, the efficacy and safety of *TUBB4A* knockout in human cells remain poorly understood. We studied patient-derived *TUBB4A^D249N^*, *TUBB4A KO*, and control individual pluripotent stem cells (iPSCs). *TUBB4A^D249N^* iPSC-derived OLs failed to mature, showing less complexity and myelination, reduced microtubule acetylation and detyrosination. *TUBB4A^D249N^* iPSC-derived MSNs also showed impaired maturation and neurite extension. *TUBB4A KO* in mutant iPSCs reduced cellular deficits and was well tolerated. These findings support that suppression of *TUBB4A* could be a safe, effective therapy for *TUBB4A*-*LD*.

## Introduction

*TUBB4A* related leukodystrophy (*TUBB4A*-LD) is a rare hypomyelinating disorder caused by autosomal dominant mutations in the *TUBB4A* gene, which encodes β-tubulin 4A. The disorder exhibits a broad spectrum of phenotypes, ranging from severe forms such as early infantile encephalopathy and late infantile hypomyelination with basal ganglia and cerebellar atrophy (H-ABC) to milder late-infantile or juvenile forms (Simons et al., 2013). A recurrent variant, p.Asp249Asn (*TUBB4A^D249N^*), accounts for over 20% of cases, especially among patients with H-ABC phenotype (Pizzino et al., 2014; Simons *et al*., 2013). H-ABC-affected patients typically develop symptoms during late infancy, including developmental delay, dystonia, ataxia, gait problems, bulbar issues, and variable cognitive deficits (Gavazzi et al., 1993; Nahhas et al., 1993). Neuroimaging features characteristic of H-ABC include hypomyelination, putamen degeneration, and cerebellar atrophy (Ferreira et al., 2014). TUBB4A is highly expressed in the cerebellum, putamen, and white matter (Hersheson et al., 2013). At a cellular level, *TUBB4A*-LD primarily affects oligodendrocytes (OLs), medium spiny neurons (MSNs) in the basal ganglia, and cerebellar granule neurons.

β-tubulins, such as TUBB4A, heterodimerize with α-tubulin to form microtubules (MTs). MTs perform critical functions, including motor protein binding, mitochondrial transport, OL process extension, myelin sheath formation, neuronal polarity, and extensions, all of which are essential for cell survival and function (Emery, 2010; Hirokawa et al., 2009; Kapitein and Hoogenraad, 2015). Using iPSC-derived models, some studies demonstrate that *TUBB4A^D249N^*variants disrupt MT dynamics and alter key cellular functions (Krajka et al., 2022; Vulinovic et al., 2018). Despite observations of these defects in *in vitro* cellular models, MSNs and OLs derived from iPSCs from individuals with H-ABC have not been published or studied to date. Our murine studies suggest that germline *Tubb4a* knockout is well tolerated and that antisense OL treatment reducing *Tubb4a* expression provides therapeutic benefit in a H-ABC mouse model (Sase et al., 2020; Sase et al., 2026). However, whether knockdown of *TUBB4A* is well-tolerated in human cells is unknown.

To model H-ABC disease mechanisms and assess whether *TUBB4A* knockdown results in phenotypic abnormalities in human cells, we generated OLs and MSNs from iPSCs derived from individuals with H-ABC harboring the recurrent *TUBB4A^D249N^* variant, as well as from *TUBB4A KO* lines. Our findings show that the *TUBB4A^D249N^* variant reduced OL and MSN differentiation and altered their complexity. Furthermore, OLs plated on nanofibers (mimicking axon-like fibers) exhibit sparse myelin sheaths and decreased myelin length. Importantly, *TUBB4A* deletion mitigates the *TUBB4A^D249N^* cellular defects observed in *TUBB4A^D249N^* variants. This is the first study to demonstrate that *TUBB4A* deletion is tolerated in iPSC-derived human neurons and OLs, supporting the potential of *TUBB4A* suppression as a treatment strategy.

## Results

### Creation of *TUBB4A* KO via germline deletion of *TUBB4A* in control and *TUBB4A^D249N^* iPSCs

To investigate the impact of the *TUBB4A^D249N^* variant in human cellular models as compared to control iPSCs (Maguire et al., 2016; Mukherjee et al., 2020; Somers et al., 2010), we generated three independent iPSC lines from fibroblasts or peripheral blood mononuclear cells (PBMCs) of individuals affected with H-ABC (Almad et al., 2023). These *TUBB4A^D249N^*iPSC lines have been thoroughly characterized previously (**Supplemental Table 1**; Almad et al., 2023). To assess the germline impact of *TUBB4A* knockout (*TUBB4A KO*), we created *TUBB4A* KO iPSC-derived cells from one of the wild-type (WT) control iPSC lines (CHOPWT6) and a KO of the *TUBB4A^D249N^* iPSC lines (LD_0638.0A) using CRISPR/Cas9 technology.

To achieve successful gene targeting, we used a dual-targeting approach with two gRNAs targeting exons 1 and 4 of *TUBB4A* on chromosome 19 in CHOPWT6 and LD_0638.0A (**Figure 1A**; **Supplemental Figure 1A)**. We refer to these lines as *TUBB4A^WT^* KO and *TUBB4A^D249N^* KO. To confirm the successful ablation of the target gene, the efficiency of the *TUBB4A* KO was validated at the mRNA level using qRT-PCR. Both the WT control iPSCs and the patient-derived *TUBB4A^D249N^* line show baseline *TUBB4A* expression. In contrast, *TUBB4A* transcript levels in both the *TUBB4A^WT^* KO and *TUBB4A^D249N^*KO lines were absent. These results demonstrate effective biallelic disruption of *TUBB4A* across both genetic backgrounds, providing a clean loss-of-function model for subsequent differentiation studies **(Figure 1B).** The clones submitted for G-band karyotyping from the *TUBB4A^WT^* KO and *TUBB4A^D249N^* KO lines showed normal karyotypes **(Figure 1C)**. The colonies that exhibited a normal karyotype were further selected to assess the pluripotency of undifferentiated iPSCs and their ability to differentiate into all three germ layers. Brightfield imaging showed compact and smooth-border colonies, typical of iPSC morphology in *TUBB4A^WT^*KO and *TUBB4A^D249N^* KO **(Figure 1D).** Undifferentiated iPSCs express pluripotency markers-OCT3/4, NANOG, and SOX2 **(Figure 1E, Supplemental Figure 1B)**. We also validated their differentiation potential by testing their ability to form the three germ layers: endoderm, mesoderm, and ectoderm. The lines successfully differentiated into these germ layers, as indicated by SOX17 for endoderm, BRACHYURY for mesoderm, and PAX6 for ectoderm (**Figure 1F**).

**Figure 1.**
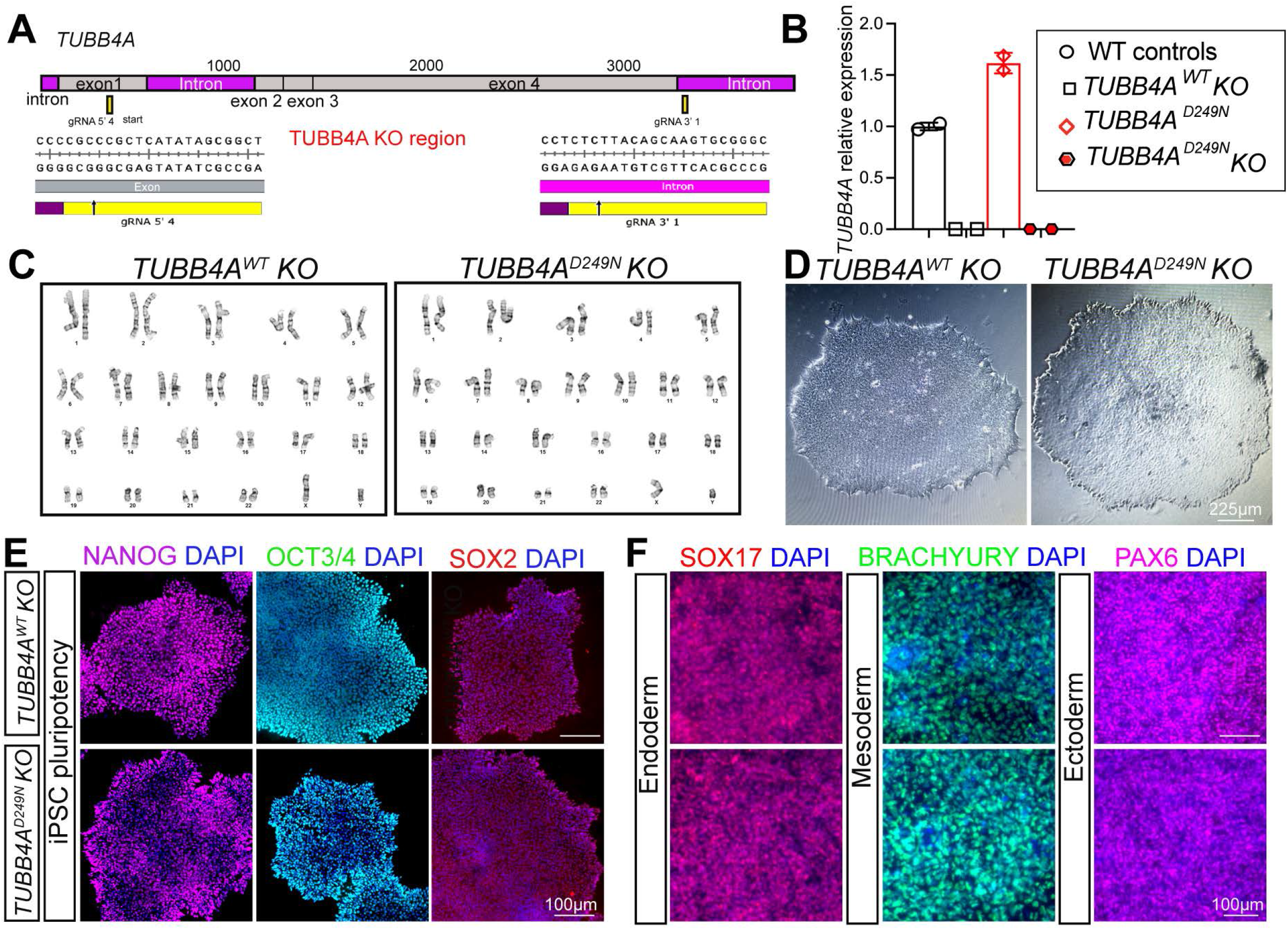
Characterization of *TUBB4A^WT^* KO and *TUBB4A^D249N^* KO iPSCs. (A) Schematic of the *TUBB4A* gene showing the position of exons, introns, and CRISPR/Cas9 gRNA target sites used for KO generation. (B) qRT-PCR analysis of *TUBB4A* mRNA levels in WT, *TUBB4A^WT^ KO* and *TUBB4A^D249N^* KO iPSCs (n=1 line per genotype and n=2-3 technical replicates per line). (C) Representative karyotype analysis of *TUBB4A^WT^* KO and *TUBB4A^D249N^* KO iPSCs, confirming normal chromosomal integrity. (D) Bright field images of *TUBB4A^WT^ KO* and *TUBB4A^D249N^* KO iPSC colonies showing typical iPSC morphology. (E) Immunofluorescence staining of pluripotency markers Nanog, OCT3/4, and SOX2 in *TUBB4A^WT^ KO* and *TUBB4A^D249N^* KO iPSCs. (F) Images showing the differentiation potential of *TUBB4A^WT^ KO* and *TUBB4A^D249N^* KO iPSCs into three germ layers: SOX17 for endoderm, BRACHYURY for mesoderm, and PAX6 for ectoderm (n=1 line per genotype and n=2-3 technical replicates per line).

### *iPSC-derived TUBB4A^D249N^* OL-like cells show poor differentiation and reduced complexity and *TUBB4A* deletion in iPSCs rescues the *TUBB4A^D249N^* OL-like phenotype

To understand the impact of *TUBB4A variants and TUBB4A* deletion in OLs and their effects on myelination, we used our validated patient-derived iPSCs with *TUBB4A^D249N^* (n=3 lines), WT control lines (n=3), *TUBB4A^WT^*KO (n=1 line, described above), and *TUBB4A^D249N^* KO (n=1 line, described above) and differentiated these iPSCs into OLs using a previously published protocol ((Douvaras and Fossati, 2015) **Figure 2A**)). Under appropriate culture conditions, control iPSCs differentiated into neural stem cells by day 8 and displayed robust Pax6 expression. By day 12, they become gliogenic cells, marked by OLIG2 (**Figure 2B**). After mechanically dissociating monolayer cells, oligospheres were generated and cultured in suspension until day 30, then replated at day 30 (**Figure 2C-D**). These plated cells were then switched to differentiating media until day 90, at which point they developed into mature OLs expressing O4+, proteolipid protein (PLP) and myelin basic protein (MBP) (**Figure 2E-I**).

**Figure 2.**
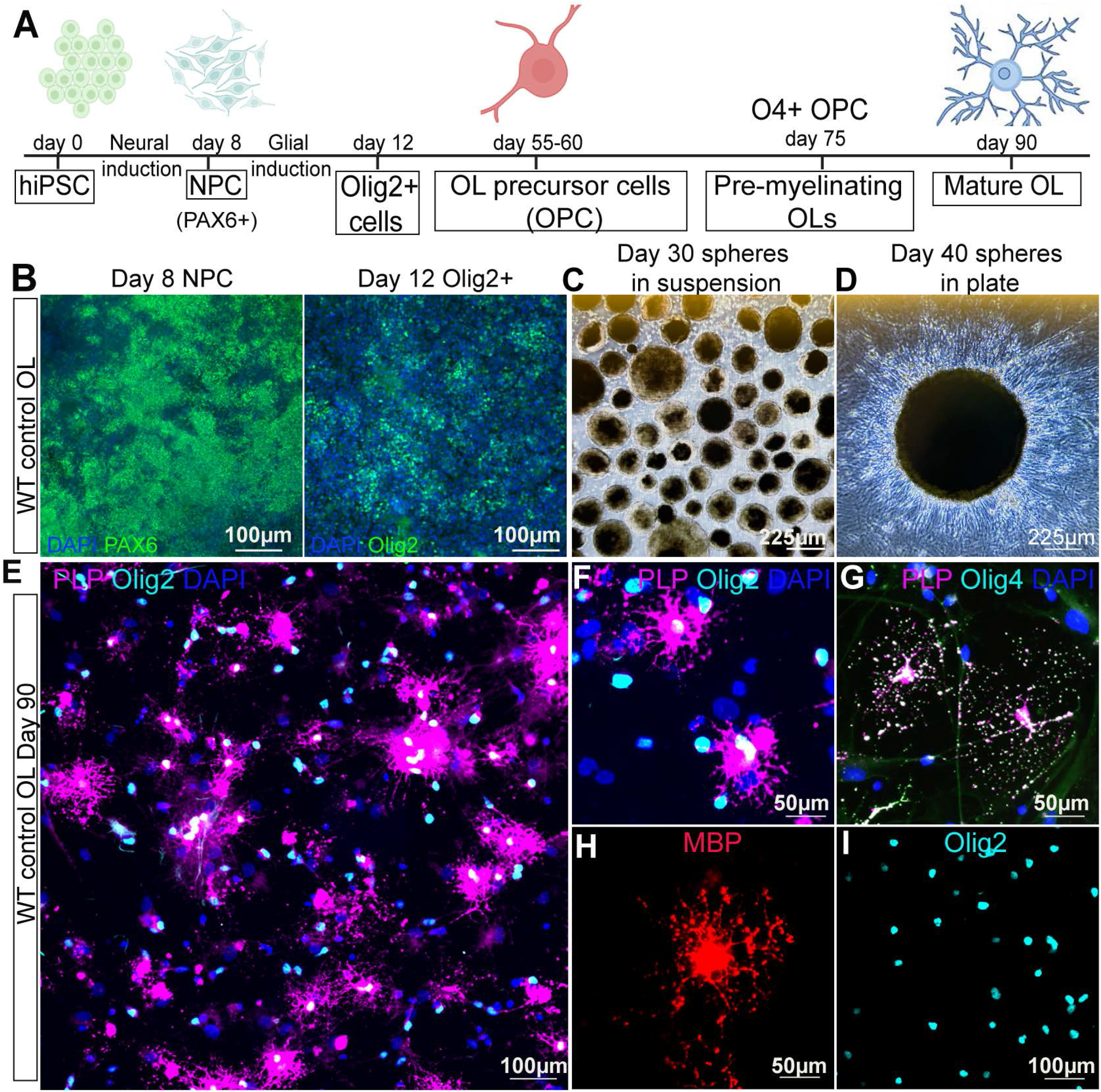
Progression and verification of iPSC-derived OL differentiation. (A) Schematic of the adapted iPSC-derived OL protocol. (B) Representative images of control iPSC-derived PAX6+ NPC at day 8 and OLIG2+ at day 12. (C) Representative image of oligospheres in suspension at day 30. (D) Representative image of plated oligospheres at day 40. (E) Representative images of OLIG2+ PLP+ WT OLs counterstained with DAPI at day 90. (F) Representative image of PLP+ OLIG2+ WT OLs at higher magnification at day 90. (G) Representative image of OLIG4+ PLP+ WT OLs counterstained with DAPI at day 90. (H) Representative image of MBP+ WT OLs at day 90. (I) Representative image of OLIG2 staining at day 90.

Next, we used this protocol to examine whether patient-derived *TUBB4A^D249N^*lines exhibit any defects in OL maturation compared to WT controls (**Figure 3A**). At day 90, we performed co-immunostaining of the cultures using PLP and OLIG2, with DAPI as the counterstain. We observed a reduced percentage of PLP+ OLs in the *TUBB4A^D249N^* lines compared to controls, indicating a possible impairment in OL differentiation (**Figure 3B and 3E**). Notably, in addition to OL differentiation defects, *TUBB4A^D249N^* OLs show distinct morphological differences compared to WT controls, appearing dystrophic, with fewer branches and reduced complexity (**Figure 3C**). We evaluated the complexity of these PLP+ OLs using Sholl analysis. *TUBB4A^D249N^* OL-like cells showed a significant decrease in the number of process intersections compared to control PLP+ OLs (**Figure 3F**). Additionally, the process lengths in *TUBB4A^D249N^* OLs appeared shorter than those in control OLs. Since MTs are key for transporting *Mbp* mRNA (Muller et al., 2013), and MBP proteins are synthesized locally on the OL processes, we assessed whether MBP protein distribution was abnormal in *TUBB4A^D249N^* OLs by immunostaining for MBP and measuring MBP density along processes. *TUBB4A^D249N^* OL processes showed reduced MBP protein on processes relative to controls (**Figure 3D and 3G**). As we noted poor MBP distribution and reduced PLP+ OLs, we assessed the *TUBB4A* OL RNA extract using a custom glia-specific gene expression panel on a custom NanoString platform. We observed significantly increased levels of gene expression for MBP, PLP, and GFAP (in a mixed culture of astrocytes, OL, and neurons) (**Supplemental Figure 3**), at odds with the ultimate protein expression. However, the *TUBB4A KO* lines, including both *TUBB4A^WT^ KO and TUBB4A^D249N^* KO, showed no abnormal phenotype and generated similar OLIG2+ cells and mature OLs as the control, with no defects in OL morphology (**Figure 3B-G and Supplemental Figure 2C**). The distribution of MBP on OL processes and the expression levels of MBP and PLP in *TUBB4A KO* OLs were consistent with those in the control lines (**Figure 3B-G and Supplemental Figure 2**).

**Figure 3.**
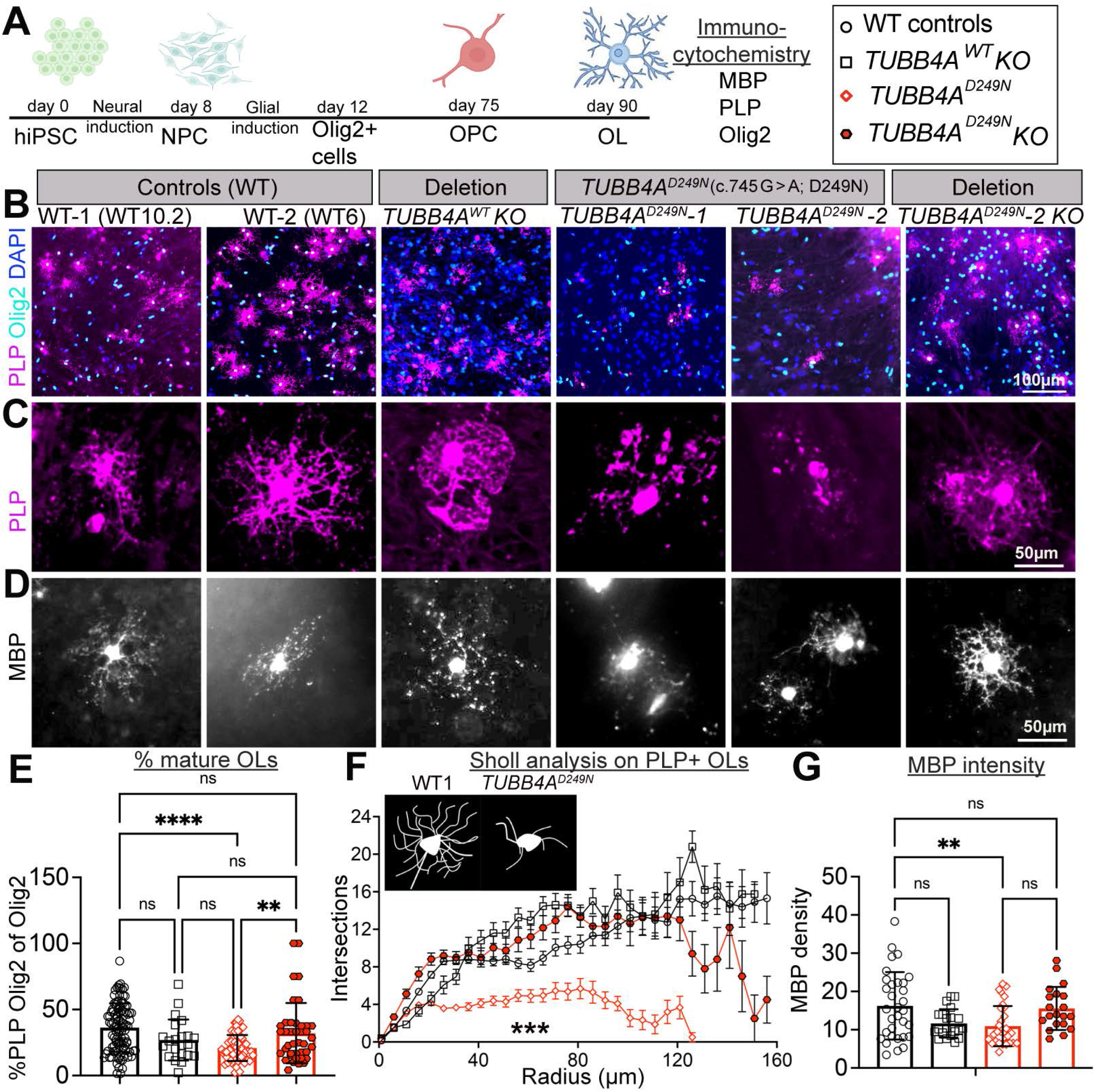
*TUBB4A^D249N^* iPSC-derived OLs fail to differentiate, show reduced complexity, while *TUBB4A KO* rescues these cellular defects. (A) Schematic of the adapted iPSC-derived OL protocol. (B) Representative images of iPSC-derived OLs stained with PLP, OLIG2, and DAPI at day 90. (C) Representative images of PLP+ OLs across different lines and genotypes. (D) Representative images of MBP+ OLs across different genotypes and lines. (E) Immunofluorescent quantification of the percentage of PLP+ OLIG2+ DAPI of OLIG2+ cells at day 90 across genotypes (WT control lines: n=3 lines; *TUBB4A^WT^KO*: n=1*; TUBB4A^D249N^*: n=3 lines; *TUBB4A^D249N^* KO: n=1; three to six technical replicates across three to four repeat differentiations). (F) Evaluation of morphological complexity in PLP+ OL-like cells using Sholl analysis software. (WT control lines: n=3 lines; *TUBB4A^WT^KO*: n=1*; TUBB4A^D249N^*: n=3 lines; *TUBB4A^D249N^* KO: n=1; n=40-60 OLs across 2-3 repeat differentiations. (G) Immunofluorescent quantification of MBP density on OL processes at D90 across genotypes (WT control lines: n=3 lines; *TUBB4A^WT^KO*: n=1; *TUBB4A^D249N^*: n=3 lines; *TUBB4A^D249N^* KO: n=1; three to six technical replicates across 2-3 repeat differentiations. *p < 0.05, **p < 0.01, ***p < 0.001. OLIG2+ graphical data and additional images are provided in Supplementary Figure 2 for each line. ns = non-significant.

### *iPSC-derived TUBB4A^D249N^*oligodendrocyte precursor cells (OPC) show abnormal morphology

Given the alteration of OL maturation in *TUBB4A^D249N^*, we next investigated whether early OPCs are affected **(Figure 4A).** On day 55, when most pre-OPCs are present based on this differentiation protocol (Douvaras and Fossati, 2015), we dissociated the oligospheres into single cells, sorted them using OPC-specific A2B5 antibody-mediated magnetic separation, and plated them on PDGF-containing medium. After 24-48 hours, the cultures were co-stained for the OPC marker A2B5 and the OL lineage marker OLIG2. Notably, sorted A2B5+ cells are multipotent; the cultures contained over 50% astrocytes and OPCs. The percentage of A2B5+ OLIG2+ out of DAPI+ cells across all lines showed no changes, indicating that the *TUBB4A* mutation does not affect OPC numbers (**Figure 4B-C**). Although the number of processes was the same between *TUBB4A^D249N^*OPCs and controls, *TUBB4A^D249N^* OPCs showed shorter processes extending from their cell bodies **(Figure 4B and 4D**). Importantly, *TUBB4A^WT^ KO* and *TUBB4A^D249N^ KO* lines exhibited no defects in OPC (**Figure 4B-E**).

**Figure 4.**
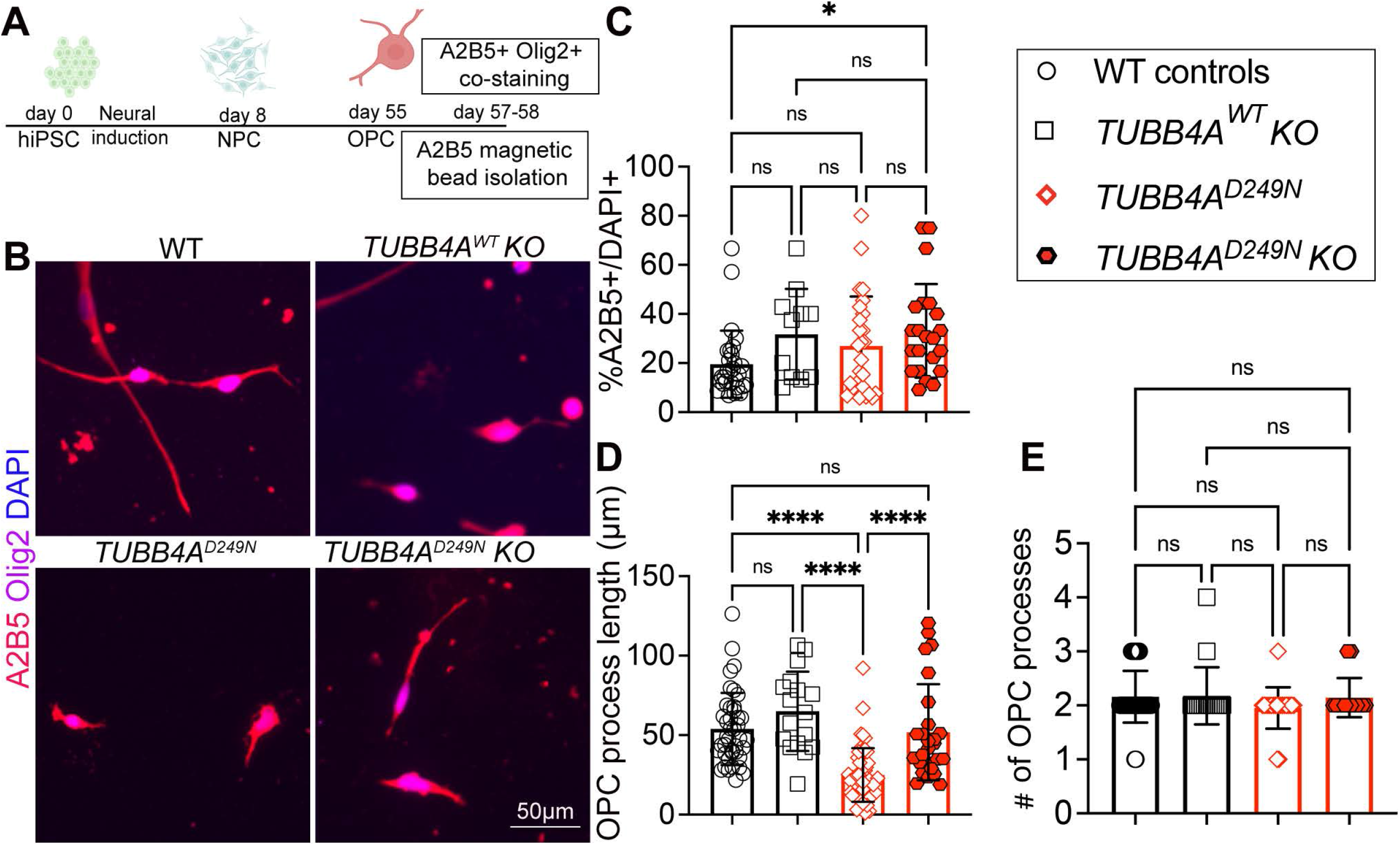
*TUBB4A^D249N^* iPSC-derived OPC exhibit reduced process length without changes in total OPC number. (A) Schematic of the adapted slow iPSC-derived OPC differentiation protocol and A2B5 magnetic isolation at day 57-58. (B) Representative images of A2B5+OLIG2+ OPCs. (C) Quantification of the percentage of A2B5+OLIG2+ of OLIG2+ cells across different genotypes. (D) Quantification of OPC process length. (E) Quantification of the number of OPC processes per cell. n=3-6 technical replicates per line with two repeat differentiations. *p < 0.05, **p < 0.01, ***p < 0.001. ns = non-significant.

### *iPSC TUBB4A^D249N^* OL shows myelin defects *in vitro* on nanofibers and reduced MT stability

To assess the myelinating potential of *TUBB4A^D249N^* OLs *in vitro*, we cultured both WT and *TUBB4A^D249N^* oligospheres on nanofibers using PDGF media from day 30 to day 75. On day 75, we switched from PDGF to Glial media to promote further maturation into myelinating cells (**Figure 5A**). Following this, we performed immunostaining for MBP and quantified the total number and length of myelin sheaths across all groups. WT control OLs effectively extended myelin sheaths on nanofibers, while *TUBB4A^D249N^*OLs displayed reductions in both the average and total sheath numbers **(Figure 5B-E)**.

**Figure 5.**
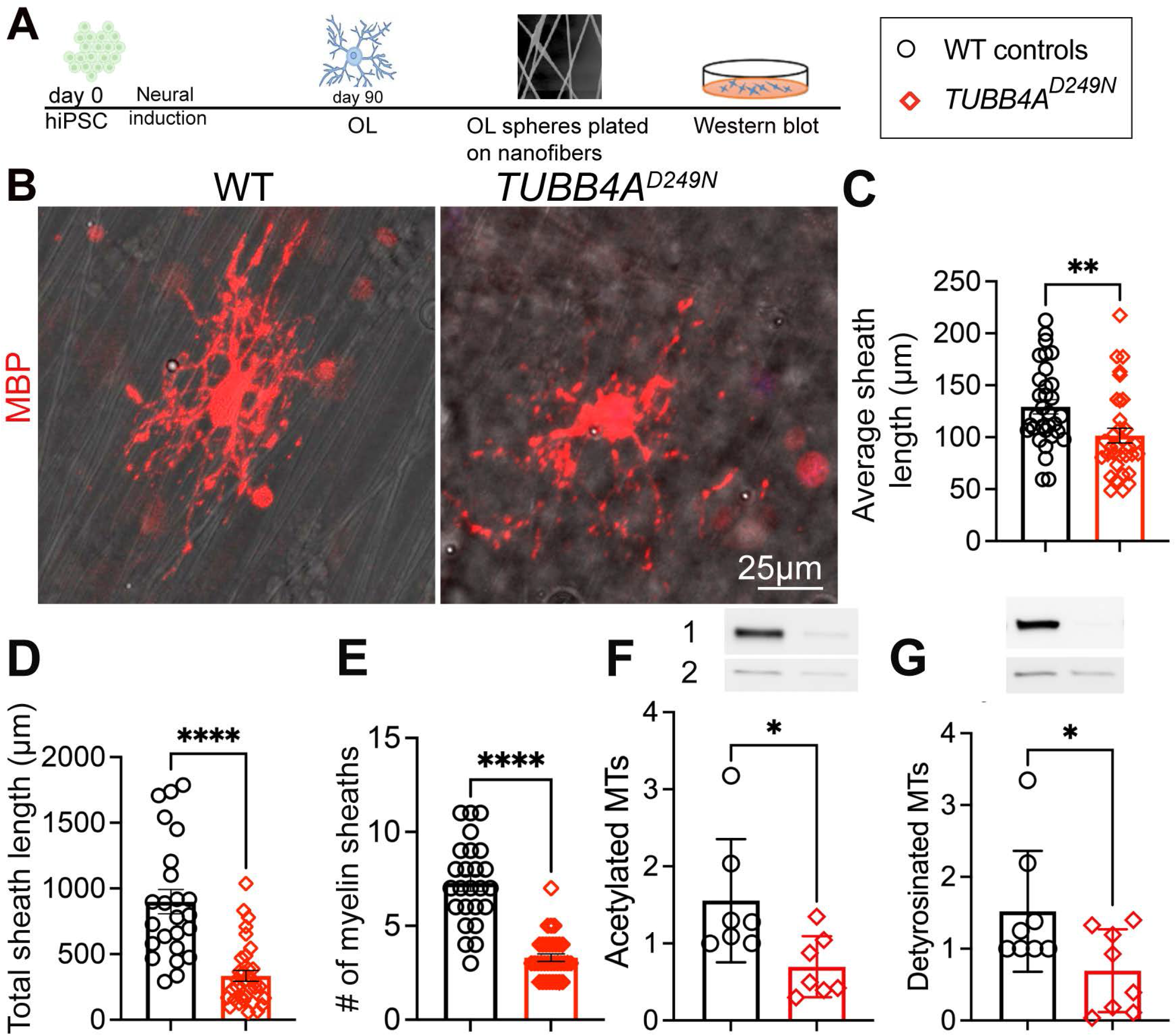
*TUBB4A^D249N^* iPSC-derived OLs myelin defects. (A) Schematic of the adapted iPSC-derived OL protocol. (B) Representative myelinating MBP+OLs on Nanofiber. (C) Quantification of Average Myelin Sheath Length (µm). (D) Quantification of the total length of myelin sheaths and (E) Number of myelin sheaths per OL. n=3 lines and 15-20 myelinating OLs with 2 repeat differentiations. (F-G) Western blot and quantification of acetylated and detyrosinated microtubules in *TUBB4A* variants versus controls. The entire set of Western blots is provided in Supplemental Figure 5, with n = 2-3 technical replicates across two repeat differentiations. *p < 0.05, **p < 0.01, ***p < 0.001.

### *iPSC TUBB4A^D249N^* OL show alterations in MT stability

To determine whether the *TUBB4A^D249N^* variant alters microtubule stability, we examined enriched protein extracts from human patient-derived iPSC-derived OLs. Microtubule stability was assessed by quantifying the post-translational modifications of α-tubulin, including acetylation and detyrosination, which are established markers of stable, long-lived microtubules. (Lu, 2025; Portran et al., 2017; Schulze et al., 1987). We measured these modifications in *TUBB4A^D249N^* and WT OLs using protein extracts from enriched OL cultures and performed Western blotting with specific antibodies. Our results reveal decreased levels of acetylated and detyrosinated MTs in *TUBB4A ^D249N^* OL **(Figure 5F and G),** indicating that the D249N variant likely reduces MT stability.

### iPSC-derived MSNs with *TUBB4A^D249N^* show maturation deficits and impaired neuronal outgrowth, while *TUBB4A* deletion rescues the phenotype

To explore striatal neuronal dysfunction, we derived MSNs from *TUBB4A* iPSCs and WT controls using a previously published protocol for MSNs (Comella-Bolla et al., 2020) (**Figure 6A**). On day 16, we derived neural progenitor cells (NPCs) from WT control lines. These cultures were verified for NPC marker expression on day 16 (PAX6+ and SOX1+) and for MSN-specific neuronal markers on day 30 (MAP2+, CTIP2+, and DARPP32+), using immunocytochemistry (**Figure 6B**).

**Figure 6.**
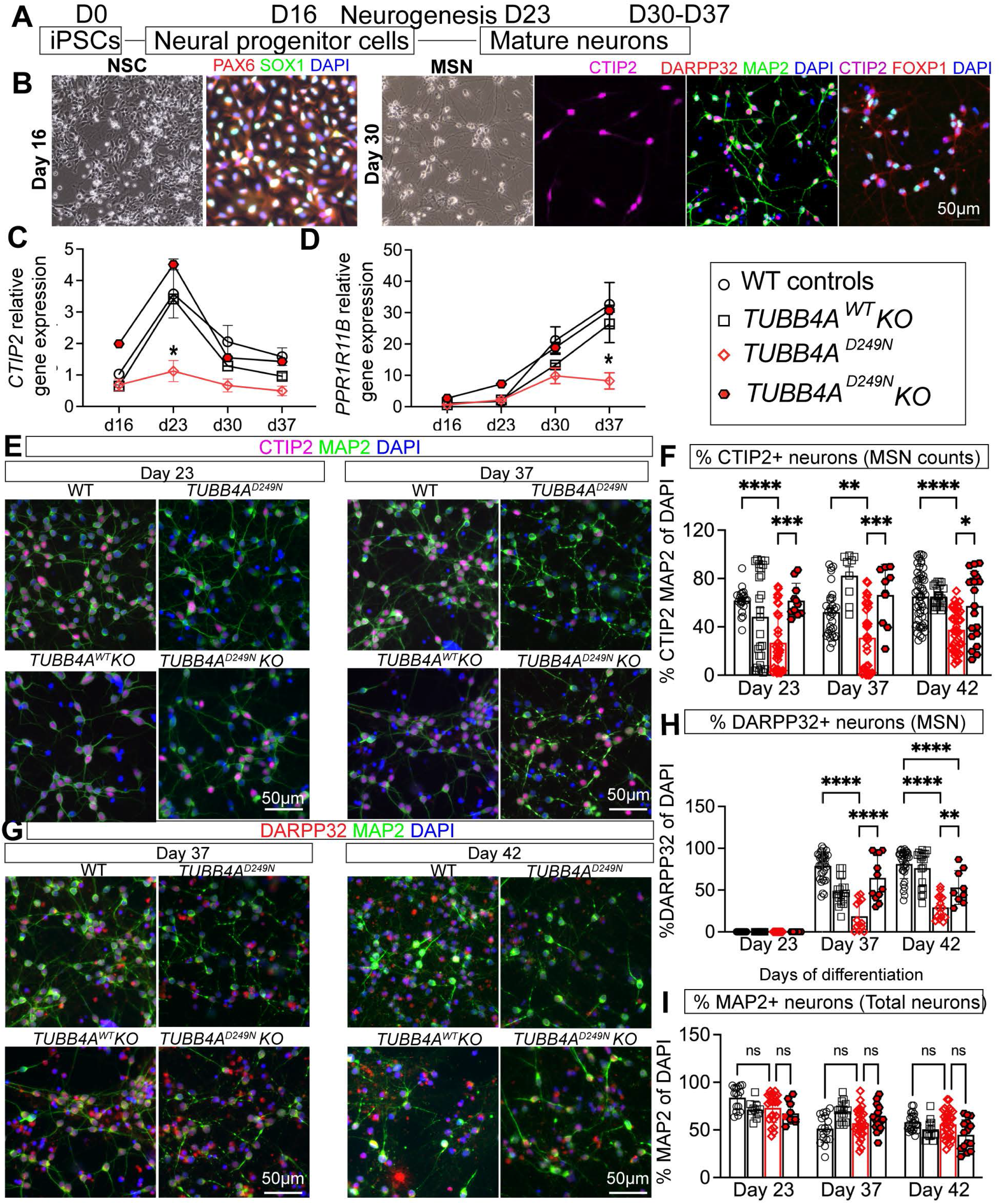
*TUBB4A^D249N^* impairs MSN mutation and neurite development, rescued by TUBB4A deletion. (A) Differentiation schematic from iPSCs to MSNs (day 0-37). (B) Representative images of NPCs at day 16 (PAX6, SOX1) and mature neurons at day 30 (CTIP2, DARPP32, MAP2), confirming MSN differentiation. (C-D) qRT-PCR of *CTIP2* and *PPP1R1B* (DARPP32) relative gene expression normalized to *HPRT* from d16 to d37 (n=3 lines per genotype and 3-6 technical replicates per genotype). (E) Representative images of CTIP2+ MAP2+ neurons with DAPI at day 23 and day 37 (F) Quantification of the total percentage of CTIP2+ MAP2+ DAPI+ neurons between all lines. (G) Representative images for DARPP32+ MAP2+ at day 37 and day 42. (H) Quantification of total percentage of DARPP32+ neurons across all lines. (I) Quantification of total percentage of MAP2+ neurons across all lines. (n=3 lines per genotype and 3-6 technical replicates per genotype with two to three repeat differentiations. *p < 0.05, **p < 0.01, ***p < 0.001. DARPP32, CTIP2, and MAP2 co-immunostaining was conducted for all time points; ImageJ was used to overlay the respective channels for Figure 6E and 6G and Supplemental Figure 4B and 4C; therefore, day 23, day 37, and day 42 image panels match for respective lines.

We then evaluated whether *TUBB4A^D249N^* lines exhibit defects in MSN development compared to WT and KO controls. Specifically, we analyzed GABAergic MSN formation by measuring the gene expression of *CTIP2* and *PPP1R1B* (DARPP32) using qPCR on days 16, 23, 30, and 37 during and after the neuronal differentiation in both control and *TUBB4A* lines (**Figure 6C**). We noted the highest CTIP2 transcript level on day 23 in control lines, although *TUBB4A^D249N^* lines showed decreased *CTIP2* expression between days 23 and 37 relative to controls (**Figure 6C**). We observed no *PPP1R1B* expression on days 16 and 23 (**Figure 6D**), consistent with the observation that CTIP2+ neurons begin expressing *PPP1R1B* after maturation (Anderson and Reiner, 1991). A decrease in *PPP1R1B* expression was noted in *TUBB4A^D249N^*lines compared to controls on days 30 and 37 (**Figure 6D**). *MAP2* gene expression remained similar across all lines (**Supplemental Figure 4A**). Next, we examined MSN defects by immunolabeling for CTIP2+, DARPP32+, and MAP2+ neurons at days 23, 30, 37, and 42. The percentage of CTIP2+ DAPI neurons was reduced in *TUBB4A^D249N^* lines compared to controls on days 23, 37, and 42 (**Figure 6E-I, Supplemental Figure 4B and 4C**). Similarly, the percentage of DARPP32+ CTIP2+ DAPI neurons decreased in *TUBB4A^D249N^*lines from days 30 to 37 (**Figure 6E-I**). The percentage of MAP2+ DAPI neurons was comparable in *TUBB4A^D249N^* versus controls at all time points (**Figure 6E, 6G and 6I**). To determine whether these neurons were undergoing cell death, we performed immunostaining for CTIP2+ and CASPASE 3+, but found no evidence of neuronal death during these periods (Data not shown). Notably, both *TUBB4A^WT^ KO* and *TUBB4A^D249N^ KO* lines exhibited comparable proportions of CTIP2+ and DARPP32+ neurons at all examined time points (**Figure 6E-I**), suggesting that the observed MSN deficits are specific to the D249N mutant and are mitigated by deletion of TUBB4A.

Given that microtubules are essential for neurite outgrowth and cytoskeletal organization (Conde and Caceres, 2009), we analyzed neurite length, soma size, and the number of primary branches. *TUBB4A* mutants displayed decreases in neurite length, primary branches, and soma size **(Figure 7A-C)**. Both *TUBB4A^WT^ KO* and *TUBB4A^D249N^ KO* lines displayed similar neurite lengths as WT controls (**Figure 7A-C**).

**Figure 7.**
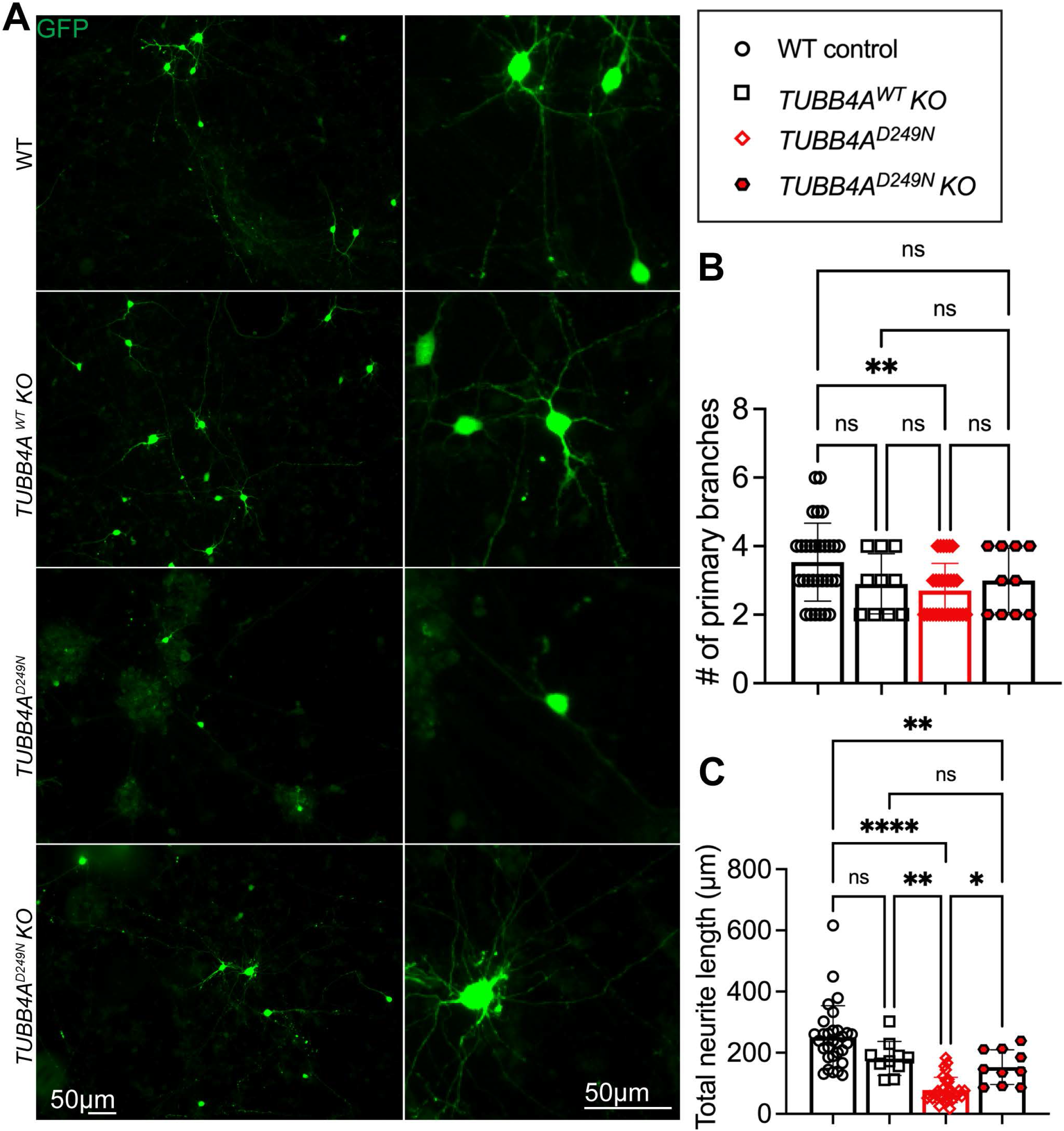
*TUBB4A^D249N^* MSN shows reduced neurite length, and deletion of *TUBB4A* does not impact the neurite length. (A) Representative images of GFP-transfected neurons showing morphologies of CTIP2+ iPSC-derived MSNs: unbranched, branched, and multipolar. (B) Quantification of the number of primary branches traced. (C) Quantification of the total neurite length traced. n=3 lines per genotype and 3-6 technical replicates per genotype. *p < 0.05, **p < 0.01, ***p < 0.001.

## Discussion

*TUBB4A-LD* is a rare neurological disorder with a spectrum of symptoms from mild to severe. The recurrent p.Asp249Asn (D249N) pathogenic variant is the most common *TUBB4*A variant and is closely associated with the radiologic findings of H-ABC, characterized by progressive myelin loss and atrophy in the striatum and cerebellum (Curiel et al., 2017; Joyal et al., 2019; Simons *et al*., 2013; van der Knaap et al., 2007; van der Knaap et al., 2002). The current study investigated disease-relevant cell types-OLs and MSN-like cells derived from *TUBB4A^D249N^* affected iPSCs. The *TUBB4A^D249N^*variant disrupts OL and neuronal differentiation, reduces cellular complexity, and impairs lineage-specific maturation. Furthermore, mutant OLs showed decreased microtubule-specific post-translational modifications-detyrosination and acetylation. Importantly, *TUBB4A* deletion restored OL and MSN maturation, supporting a model in which *TUBB4A* dysfunction causes cell-autonomous toxic gain-of-function effects in vulnerable cell types, thereby contributing to hypomyelination and striatal pathology in H-ABC.

### Maturation and Morphology

When we differentiated our *TUBB4A^D249N^* iPSCs into OPCs and compared them with control iPSC-derived OPCs, we found the same number of OPCs and processes, but shorter outgrowths compared with controls. Because *TUBB4A* expression is relatively low in OPCs compared with differentiating OLs, we did not anticipate OPC deficits *in vitro* (Hersheson *et al*., 2013). Since these OPCs are mitotic and proliferate, the number of OPCs in mutants was unchanged; therefore, we did not directly assess OPC proliferation or cell death in vitro. However, our *Tubb4a^D249N^* mouse model showed increased OPC proliferation secondary to OPC death, resulting in unchanged overall OPC numbers (Sase *et al*., 2020). Consistent with this possibility, *TUBB4A^D249N^*OL-like cells show reduced PLP+ OLs. The reduced number of OLs in the *TUBB4A^D249N^* variant might result from abnormal OPC differentiation or increased susceptibility of newly formed OLs to apoptosis, or a combination of both factors (Emery and Wood, 2024; Gacem and Nait-Oumesmar, 2021). Furthermore, *TUBB4A^D249N^* OL-like cells exhibited impaired process extension, branching, and poor MBP distribution. These findings are consistent with the established role of microtubules in OL cytoskeletal formation and with impaired intracellular transport in rodent *TUBB4A* models (Duncan et al., 2017; Krajka *et al*., 2022; O’Connor et al., 2000). Despite reduced MBP protein distribution in OLs, our data indicate increased *MBP* transcript levels in mutant OLs relative to controls, suggesting a compensatory transcriptional response to defective protein trafficking or localization (Buccitelli and Selbach, 2020). In addition, our data show that enriched *TUBB4A^D249N^* OL-like cells exhibit reduced microtubule-specific acetylation and de-tyrosination, reflecting alterations in post-translational modifications. The mechanisms by which the D249N variant alters microtubule post-translational modifications remain unclear, but previous studies have shown that the D249N variant alters microtubule dynamics, tubulin conformations, and organization, which may contribute to altered post-translational modification (Krajka *et al*., 2022; Sase *et al*., 2020). Overall, these findings suggest that the D249N variant disrupts mitotic OPCs and post-mitotic OLs. Future research is needed to explore whether OPC abnormalities directly lead to failures in OL maturation and myelination or are separate events.

### Myelination and Microtubule Stability

To assess the ability of OLs to myelinate, we differentiated wild-type and *TUBB4A^D249N^* OL-like cells on nanofibers. *TUBB4A* mutant OL-like cells generated fewer myelin sheaths and significantly shorter myelin sheaths than controls, suggesting defective myelin initiation and elongation (Fu et al., 2019). Essentially, the myelin sheath contains the microtubules known as lamellar microtubules (Fu *et al*., 2019). These microtubules are within cytoplasmic channels called myelinic channels, which extend through the myelin sheath, and likely facilitate vesicular transport during myelination (Edgar et al., 2021; Kassmann et al., 2011; Weigel et al., 2021). Previous studies have demonstrated that TPPP, a microtubule nucleation factor that is predominantly expressed in OLs, is required for myelin elongation and that its loss results in shortened lamellar microtubules and impaired sheath elongation but does not affect myelin initiation (Fu *et al*., 2019). Given the reduced myelin sheath initiation and elongation in *TUBB4A^D249N^* OLs, it is possible that this variant disrupts microtubule organization and stability within developing myelin sheaths. This could impair the transport of essential myelin proteins, such as MBP, and other myelin components (lipids) and result in dysregulation of myelin sheath development and maintenance.

In addition to oligodendrocyte abnormalities, we show that *TUBB4A^D249N^* MSN-like cells displayed impaired differentiation and reduced neurite outgrowth. An important unresolved question is how microtubule dysfunction reduces the number of CTIP2+ and DARPP32+ neurons. CTIP2 maturation is dependent on BDNF–TrkB signaling (Braid and Johnson, 1992; Rauskolb et al., 2010), and BDNF-containing vesicles are transported along microtubules (Gauthier et al., 2004). Therefore, impaired microtubule dynamics may disrupt BDNF vesicular trafficking, thereby compromising MSN maturation and development. Future studies examining neurotrophic factor transport and signaling in *TUBB4A*-mutant neurons may provide mechanistic insights into striatal pathology.

Although the role of TUBB4A in myelination and neuronal differentiation remains poorly understood, emerging molecular therapies are promising. Our recent preclinical studies in the H-ABC mouse model showed that reducing *Tubb4a* with antisense oligonucleotides rescued phenotypes *in vivo* (Sase *et al*., 2026). While *Tubb4a* knockdown appears safe in mice, its safety in human cells was unclear. We created *TUBB4A KO* iPSCs from control and wild-type cells and differentiated them into OL-like and MSN-like cells. Germline *TUBB4A* deletion was well tolerated, with no differentiation issues or morphological deficits in OL and MSN-like cells. These results support the safety of *TUBB4A* reduction, but because tubulins are essential during development, assessing whether knockdown affects stem cell differentiation or disturbs the microtubule dynamics is necessary. Together, iPSC-derived OL-and MSN-like cells are normal and able to generate OLs and MSN-like cells in the absence of *TUBB4A*. Overall, our data support evidence that reducing *TUBB4A* expression may represent a promising therapeutic strategy for TUBB4A-associated leukodystrophy. These findings provide evidence of cell-autonomous deficits associated with *TUBB4A* variants and preclinical support for the continued development of TUBB4A-targeted downregulation therapies, as well as ongoing efforts to translate this approach toward clinical application.

## Methods

### iPSC Generation, Characterization, and Maintenance

*TUBB4A* iPSC lines used in this study include control, patient-derived, and gene-edited knockout lines (**see Supplemental Table 1**). Three control iPSC lines—CHOPWT4, CHOPWT6, and CHOPWT10.2—were obtained from the Children’s Hospital of Philadelphia (CHOP) iPSC Core (Maguire *et al*., 2016; Mukherjee *et al*., 2020; Somers *et al*., 2010). Two homozygous *TUBB4A* knockout lines were created using CRISPR-Cas9: CHOPWT6 KO (clone 6), derived from the CHOPWT6 control line, and LD_0638.0A KO (clone 39), derived from the patient line LD_0638.0A. Patient-derived lines LD_0313.0, LD_0638.0A, and LD_0853.0 carry the heterozygous *TUBB4A* D249N mutation and are well-characterized (Almad et al., 2023). All lines were maintained in feeder-free conditions on Matrigel-coated plates using mTeSR Plus medium (Stem Cell Technology) with daily media changes. After recovery from thawing, each line was passaged two to three times before use. Pluripotency was confirmed through trilineage differentiation.

### Biological materials and ethical considerations

All methods related to human iPSC culture and differentiation received ethical approval from the Office for IRB (IRB 14-011236) and were performed in accordance with the relevant institutional guidelines and regulations, including the Declaration of Helsinki and the Ethical Guidelines for Medical and Health Research Involving Human Subjects.

### Differentiation towards Medium Spiny Neurons

Human iPSCs were differentiated into medium spiny neurons using an adapted protocol based on (Comella-Bolla *et al*., 2020). Briefly, neural induction was achieved through dual inhibition of SMAD and WNT signaling, followed by patterning of ventral telencephalic progenitors. Neuronal differentiation was initiated at day 16 using differentiation medium containing 1:1 (Advanced DMEM/F12 (Gibco) and Neurobasal medium (Gibco) supplemented with 2 mM GlutaMax (Gibco), 1X B27 with RA (Gibco), 2 μM PD0332991 (Tocris), 10 μM DAPT (Tocris), 10 ng/ml BDNF (R&D Systems), 10 μM Forskolin (Tocris), 3 μM CHIR (Tocris), 300 μM GABA (Tocris), 1.8 mM CaCl2 (Sigma) and 200 μM Ascorbic acid (Sigma)) and cells matured into MSNs over several weeks using differentiation medium containing 1:1 (Advanced DMEM/F12 (Gibco) and Neurobasal medium (Gibco) supplemented with, 2 mM GlutaMax (Gibco), 1X B27 with RA (Gibco), 2 μM PD0332991 (Tocris), 10 μM DAPT (Tocris), 10 ng/ml BDNF (R&D Systems), 10 μM Forskolin (Tocris), 3 μM CHIR (Tocris), 300 μM GABA (Tocris), 1.8 mM CaCl2 (Sigma) and 200 μM Ascorbic acid (Sigma)). Cells were harvested at key time points (days 16, 23, 37 and 42) for downstream analysis.

### Differentiation towards Oligodendrocytes

A modified version of the protocol from (Douvaras and Fossati, 2015) was used to differentiate iPSCs into oligodendrocyte-like (OL-like) cells. iPSCs were first converted to PAX6^⁺^ neural progenitors and then to OLIG2^⁺^ progenitors, which formed spheres in suspension within low-attachment 6-well plates at day 12. To improve differentiation efficiency, spheres were cultured on an orbital shaker inside the incubator (88 rpm, 19 mm orbit) from days 12 to 20 into N2B27 media (DMEM/F12 (Gibco), 1X NEAA (Gibco), 1X GlutaMax (Gibco), 1X Penicillin/Streptomycin (Gibco), 1X Beta-mercaptoethanol (MP Biomedicals), 1X N2 (Gibco), 1X B27-(w/o VitA) (Gibco), 25 μg/ml Insulin (Sigma), 100 nM RA (Sigma), and 1 μM SAG (Tocris)) and then at day 20 transferred into PDGF media (DMEM/F12 (Gibco), 1X NEAA (Gibco), 1X GlutaMax (Gibco), 1X Penicillin/Streptomycin (Gibco), 1X Beta-mercaptoethanol (MP Biomedicals), 1X N2 (Gibco), 1X B27-(w/o VitA) (Gibco), 10 ng/mL PDGF-AA (PeproTech), 5 ng/mL HGF (R&D Systems), 10 ng/mL IGF-1 (R&D Systems), 10 ng/mL NT3 (PeproTech), 100 ng/mL Biotin (Sigma), 60 ng/mL T3 (Sigma), 1 μM cAMP (Sigma), and 25 μg/mL insulin (Sigma)). At day 30, spheres were plated in PDGF media onto 24-well plates containing glass coverslips and 6-well plates coated with poly-L-ornithine and laminin, following the original plating methods. Terminal differentiation in the glial medium (DMEM/F12 (Gibco), 1X NEAA (Gibco), 1X GlutaMax (Gibco), 1X Penicillin/Streptomycin (Gibco), 1X Beta-mercaptoethanol (MP Biomedicals), 1X N2 (Gibco), 1X B27-(w/o VitA) (Gibco), 10 mM HEPES (Gibco), 5 ng/mL HGF (R&D Systems), 10 ng/mL IGF-1 (R&D Systems), 10 ng/mL NT3 (PeproTech), 100 ng/mL Biotin (Sigma), 60 ng/mL T3 (Sigma), 1 μM cAMP (Sigma), 25 μg/mL insulin (Sigma) and 20 μg/mL Ascorbic Acid (Sigma) was performed for 14 days. All assays, including immunocytochemistry, RNA, and protein extraction for mature OLs, were conducted on mixed glial cultures. All reagent information is provided in Supplemental Table 3.

### Oligosphere Dissociation

Oligospheres were dissociated into single cells at differentiation day 55. Papain (Thermo scientific) was activated in enzyme activation buffer containing EDTA (Thermo scientific), L-cysteine HCl (Sigma), and β-mercaptoethanol (Gibco) and incubated at 37°C for 30 min. Ovomucoid solution (Sigma) and all dissociation buffers were prepared in DPBS () and warmed to RT prior to use. Cultures were washed once with DMEM/F12 and incubated with activated papain at 37°C for 15 min. Cells were gently dissociated by pipetting to detach neurospheres and transferred to 15 mL tubes. Enzymatic digestion was stopped by adding ovomucoid solution at a 1:1 ratio. The supernatant containing single cells was collected and placed on ice. Residual spheres were further digested using Enzyme P, Buffer Y, and Enzyme A according to the manufacturer’s instructions (Miltenyi Biotec), followed by incubation at 37°C on rotator. Mechanical dissociation was performed using sequential pipetting with large-, medium-, and small-bore glass pipettes to generate a single-cell suspension, which was then combined with the previously collected single-cell fraction. Cell suspensions were filtered through a pre-wetted 70 µm separation filter, washed with PDGF-containing medium, centrifuged at 400 × g for 5 min, and counted.

### A2B5 Magnetic-Activated Cell Sorting (MACS)

After dissociation of oligospheres, dissociated cells were resuspended in MACS buffer and incubated with FcR receptor blocking reagent (Miltenyi Biotec, 130-059-901) for 10 min at 4°C, followed by incubation with A2B5 antibody (Miltenyi Biotec, 130-093-392) according to the manufacturer’s protocol. Cells were washed, centrifuged at 300 × g for 10 min, and applied to pre-equilibrated MACS columns. Columns were washed with MACS buffer, and A2B5-positive cells were eluted, counted, and centrifuged at 300 × g for 5 min prior to downstream applications.

### Plating on nanofibers to study myelination potential

To assess the myelination potential of iPSC-derived OL, the neurospheres were plated onto aligned nanofiber scaffolds (Sigma, Z694614) at day 30. Nanofibers were attached to tissue culture plates using a sterile silicone gel (Kwik-Cast, WPI Inc.) and coated with poly-L-ornithine (Sigma, P3655) and laminin (Sigma, L2020). The spheres were plated, and the cells were allowed to mature on the nanofibers for 90 days in PDGF media. At that point, they were fixed and stained via immunocytochemistry (ICC) with respective myelin markers.

### Immunocytochemistry

Coverslips from iPSC, MSN, and OL-mixed cultures were fixed in 4% paraformaldehyde (PFA) for 15 minutes at room temperature, followed by three washes with PBS. Samples were then blocked and permeabilized in 5% normal goat serum (Sigma) and 0.2% Triton X-100 (Sigma) for 1 hour at room temperature. Coverslips were incubated overnight at 4 °C with primary antibodies **(Supplemental Table 2)**. For MSNs, on day 23, day 37, and day 42, co-staining for CTIP2, MAP2, and DARPP32 was conducted. For OLs, on day 90, co-staining for PLP, Olig2 and MBP was conducted. The next day, cells were washed three times with PBS and incubated with appropriate fluorescently labeled secondary antibodies for 30 minutes at room temperature. After an additional three PBS washes, coverslips were mounted onto glass slides using ProLong™ Gold Antifade Mountant with DAPI (Thermo Fisher Scientific). Images were taken using Keyence and Confocal microscopy with a 20× objective for cell quantification and a 40× objective for Sholl analysis, branch count, and neurite length measurements. For quantification, n=3–4 replicate wells were analyzed per condition for each iPSC line.

### Immunoblotting

After differentiation into mature OL-like cells, enriched OLs cultures were lysed in RIPA buffer (Thermo Fisher Scientific) supplemented with protease and phosphatase inhibitor cocktails (Sigma-Aldrich). Protein lysates were diluted with 4× Laemmli sample buffer (Bio-Rad), boiled for 5 min in 95°C, and electrophoresed on SDS-PAGE using Any kD™ Mini-PROTEAN® TGX™ Precast Protein Gels (Bio-Rad, Hercules, CA, USA). Proteins were transferred onto 0.45-µm LF PVDF membranes (Trans-Blot Turbo RTA Mini 0.45 µm LF PVDF Transfer Kit , Bio-Rad) using the Trans-Blot® Turbo Transfer System (Bio-Rad). Membranes were blocked for 1 h at room temperature in 1% non-fat dry milk (Bio-Rad) prepared in Tris-buffered saline containing 0.05% Tween-20 (TBST) and incubated overnight at 4°C with rabbit anti-detyrosinated α-tubulin (1:1,000; RevMAb, Cat. No. 31-1335-00) diluted in blocking buffer. The membranes were washed 5 times with TBST and incubated for 1 h at room temperature with IRDye® 800CW Donkey Anti-Rabbit IgG secondary antibody (1:5,000; LI-COR Biosciences) diluted in blocking buffer. After washing with TBST, fluorescent signals were detected using an Odyssey Infrared Imaging System (LI-COR Biosciences) at 800 nm. The membranes were then re-probed with mouse anti-acetylated tubulin (1:1,000; Sigma-Aldrich, Cat. No. T7451), followed by incubation with IRDye® 680RD Donkey Anti-Mouse IgG secondary antibody (1:5,000; LI-COR Biosciences), and imaged at 700 nm using the Odyssey system, or vice versa. We cross-verified and confirmed that there is no signal interference (Data not shown). Images were analyzed using FIJI software. For normalization with loading control, membranes were stripped according to the manufacturer’s instructions using stripping buffer (Restore™ Fluorescent Western Blot Stripping Buffer, Thermo) and re-probed with mouse anti-GAPDH (1:3,000; Sigma, Cat. No. MAB378) or mouse anti-vinculin (1:3,000; Sigma-Aldrich, Cat. No. V9131).

### RNA Extraction, qRT-PCR and Nanostring

Total RNA was isolated from both MSN and mixed OL cultures in 6-well plates using the RNeasy Micro Kit (Qiagen, 74004) including the optional 15-minute DNAse incubation per manufacturer’s instructions. cDNA was generated from 200 ng of RNA per sample using the High-Capacity RNA-to-cDNA™ Kit (Thermo Fisher Scientific, 4388950). Gene expression was analyzed through qRT-PCR using the Applied Biosystems QuantStudio 7 Flex (Thermo Fisher Scientific). Relative expression was calculated using the 2^-ΔΔCt^ method with normalization to GAPDH. Prime Time qPCR Assay *TUBB4A* and *GAPDH* and/or *HPRT* (Integrated DNA Technologies, Hs00258900_m1, Hs00259967_m1 and Hs01102259_m1 respectively). Replicates of this solution were placed in a 384-well plate for QuantStudio FLEX 7 qRT-qPCR (Thermo Fisher). Enriched RNA from oligospheres at day 90, 500ng, were hybridized with the nCounter CodeSets (nanoString, Seattle, USA) to a custom-designed human Nanostring panel, prepped on nCounter FLEX prep station (nanoString) and scanned the image on nCounter FLEX digital analyzer (nanoString). Raw counts were normalized to the internal positive controls and 4 housekeeping genes (*ALAS1, HPRT, PPIB and TBP*) by following the company’s instructions.

### Transfection of MSNs

For neurite length analysis, we transfected iPSC-derived MSNs on day 37 using Lipofectamine Stem Transfection Reagent (Thermo Fisher) and 1.25 μg plasmid DNA (PGK-EGFP (Addgene #216110)) per dish as per the manufacturer’s instructions. After 24-48 h of incubation, cells were fixed and co-stained with the CTIP2 marker for identification. The GFP-transfected neurons were imaged and analyzed as described in the Image analysis.

### Image analysis

#### MSN and OL cell count quantification

Cell quantification was performed using the Keyence Analyzer software and/or Fiji (ImageJ). Individual cells positive for the markers of interest were automatically and manually identified and counted in each image.

#### MSN and OL complexity measurements

MSN and OL complexity was quantified using Sholl analysis and neurite tracing. Sholl analysis was conducted on PGK-GFP-transfected MSNs and PLP-stained OLs, counting the intersections of neuronal or OL processes with a series of evenly spaced concentric circles originating from the cell body as conducted previously (REFF). The Sholl analysis function is found within the SNT plugin of Fiji’s *Neuroanatomy* update site. Neurite tracing was performed manually, and neurite length and branch number were measured using the NeuronJ plugin in Fiji. For both Sholl analysis and neurite tracing, only MSNs and OLs that were distinguishable from neighboring cells were used for analysis.

#### Myelin sheath assessments of nanofibers

Quantitative image analysis of myelin sheath assessments was performed using Fiji ImageJ. OL processes associated with and wrapping around nanofibers were analyzed using the NeuronJ plugin. Individual myelinating processes were manually traced along the length of the nanofibers to measure process length. Myelin sheath complexity was quantified by counting branch points per traced process. All measurements were conducted using identical analysis parameters across experimental conditions.

## Statistical analysis

Each differentiation experiment was repeated at least two to three times. The number of technical replicates included in each experiment is listed in the figure legends. For statistical analysis, one-way or two-way ANOVA, and t-test with normal distribution, or one-way Chi-square tests were used as appropriate. * Indicates p value < 0.05, ** p value < 0.01, *** p value < 0.001, n.s. = not significant. Data are presented as mean ± SEM and detailed biological and technical replicates are provided in the figure legends

**Supplemental Figure 1. Validation and Characterization of *TUBB4A* KO iPSC Lines** (A) Schematic representation of the CRISPR/Cas9 strategy for *TUBB4A* gene disruption. (B) Characterization of pluripotency in *TUBB4A^WT^* KO and *TUBB4A^D249N^* KO clones. (C) Expected PCR product sizes. (D) PCR validation of *TUBB4A* KO clones.

**Supplemental Figure 2.** (A and B) Representative images of iPSC-derived OLs stained with PLP, OLIG2, and DAPI at day 90 across different lines of *TUBB4A* and controls. (C) Immunofluorescent quantification of the percentage of OLIG2+ cells of DAPI cells at day 90 across genotypes (*TUBB4A^D249N^*: n=3 lines; *TUBB4A^D249N^* KO: n=1; three to six technical replicates across three to four repeat differentiations). *p < 0.05, ***p < 0.01, ****p < 0.001. ns = non-significant.

**Supplemental Figure 3** RNA expression levels in enriched OL cultures at day 90 measured using a custom NanoString across different genotypes (n=3-9 technical replicates across 2 differentiations). ns= non-significant

**Supplemental Figure 4:** (A) qRT-PCR of *MAP2* relative gene expression normalized to *HPRT* from d16 to d37 (n=3 lines per genotype and 1-3 technical replicates per genotype). (B) Representative images of CTIP2+ neurons at day 42. (C) DARPP32+ neurons (No DARPP32 is detected at day 23). Quantification is provided in Figure 6. DARPP32, CTIP2, and MAP2 co-immunostaining was conducted for all time points; ImageJ was used to overlay the respective channels for Figure 6E and 6G and Supplemental Figure 4B and 4C; therefore, day 23, day 37, and day 42 image panels match for respective lines.

**Supplemental Figure 5:** (A–C) Immunoblotting images showing bands for acetylated MTs and respective GAPDH or vinculin blots. (D–F) Immunoblotting images showing bands for de-tyrosinated MTs and respective GAPDH blots. For Runs 1–2, the membrane was first probed, imaged for the first protein, re-probed for the second protein with a different host species and different emission spectra (MT acetylation – 680nm; MT detyrosination – 800nm), and subsequently stripped to obtain the GAPDH bands.

**Supplementary Table 1:** iPSC-derived TUBB4A lines

**Supplementary Table 2:** Antibody resource Table

## Resource availability

### Lead contact

Adeline Vanderver.

### Materials availability

This study did not produce any new unique reagents. Unless noted otherwise, data are available in the main text or supplementary materials.

### Data and code availability

## Acknowledgements

We would like to thank Dr. Judith B. Grinspan for providing hybridoma antibodies. This study was partly supported by funds from Synaptix Bio company; Kamens Chair in Translational Neurotherapeutics; H-ABC Foundation UK; and Commonwealth Universal Research Enhancement Program (CURE) funding (SAP # 4100077047).

## Author contributions

Conceptualization – SS, AA, and AV; Methodology -ZH, DB, PRN, SS, LG, TJ, JLH, AB, AA, AT Software-ZH, SS, DB, LG, AA; Formal analysis-ZH, SS, DB, PRN; Investigation –SS, ZH, AV; resources - AV; Writing (original draft) – ZH, SS and DB; Writing – review and editing ZH, SS, AV, JG, DB, AA, LG, JLH; Visualization –SS and AV; Supervision – SS, AA and AV; Funding acquisition - AV

## Declaration of generative AI and AI-assisted technologies in the writing process

During editing, the authors used Grammarly to review the grammar. All minor adjustments were made by the authors, and we accept full responsibility for the publication’s content.

