## Supplemental data for "Deletion of *TUBB4A* mitigates the oligodendrocyte and neuronal deficits in human iPSCs derived from individuals affected by H-ABC"

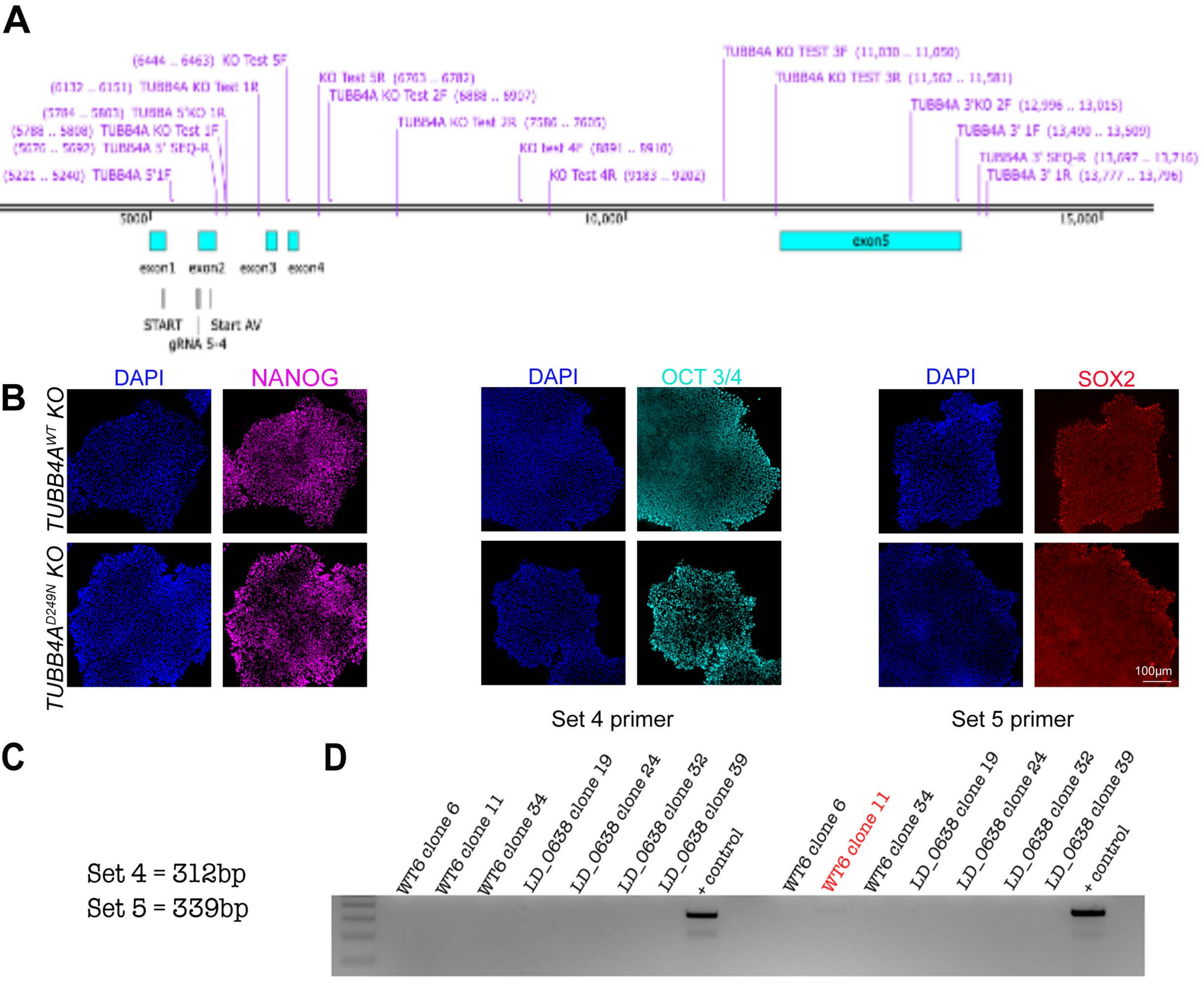
A
(6444 .. 6463) KO Test 5F
(6132 .. 6151) TUBB4A KO Test 1R
(5784 .. 5803) TUBBA 5'KO 1R
(5788 .. 5808) TUBB4A KO Test 1F
(5221 .. 5240) TUBB4A 5'1F
KO Test 5R (6763 .. 6782)
TUBB4A KO Test 2F (6888 .. 6907)
TUBB4A KO Test 2R (7586 .. 7606)
KO Test 4R (9183 .. 9202)
TUBB4A KO TEST 3F (11,030 .. 11,050)
TUBB4A KO TEST 3R (11,562 .. 11,581)
TUBB4A 3'KO 2F (12,996 .. 13,015)
TUBB4A 3' 1F (13,490 .. 13,509)
TUBB4A 3' SEQ-R (13,697 .. 13,716)
TUBB4A 3' 1R (13,777 .. 13,796)
5000
10,000
15,000
exon1
exon2
exon3
exon4
exon5
START
Start AV
gRNA 5-4
B
DAPI
NANOG
DAPI
OCT 3/4
DAPI
SOX2
TUBB4A^WT KO
TUBB4A^D249N KO
100μm
Set 4 primer
Set 5 primer
C
Set 4 = 312bp
Set 5 = 339bp
D
WT6 clone 6
WT6 clone 11
WT6 clone 34
LD_0638 clone 19
LD_0638 clone 24
LD_0638 clone 32
LD_0638 clone 39
+ control
WT6 clone 6
WT6 clone 11
WT6 clone 34
LD_0638 clone 19
LD_0638 clone 24
LD_0638 clone 32
LD_0638 clone 39
+ control

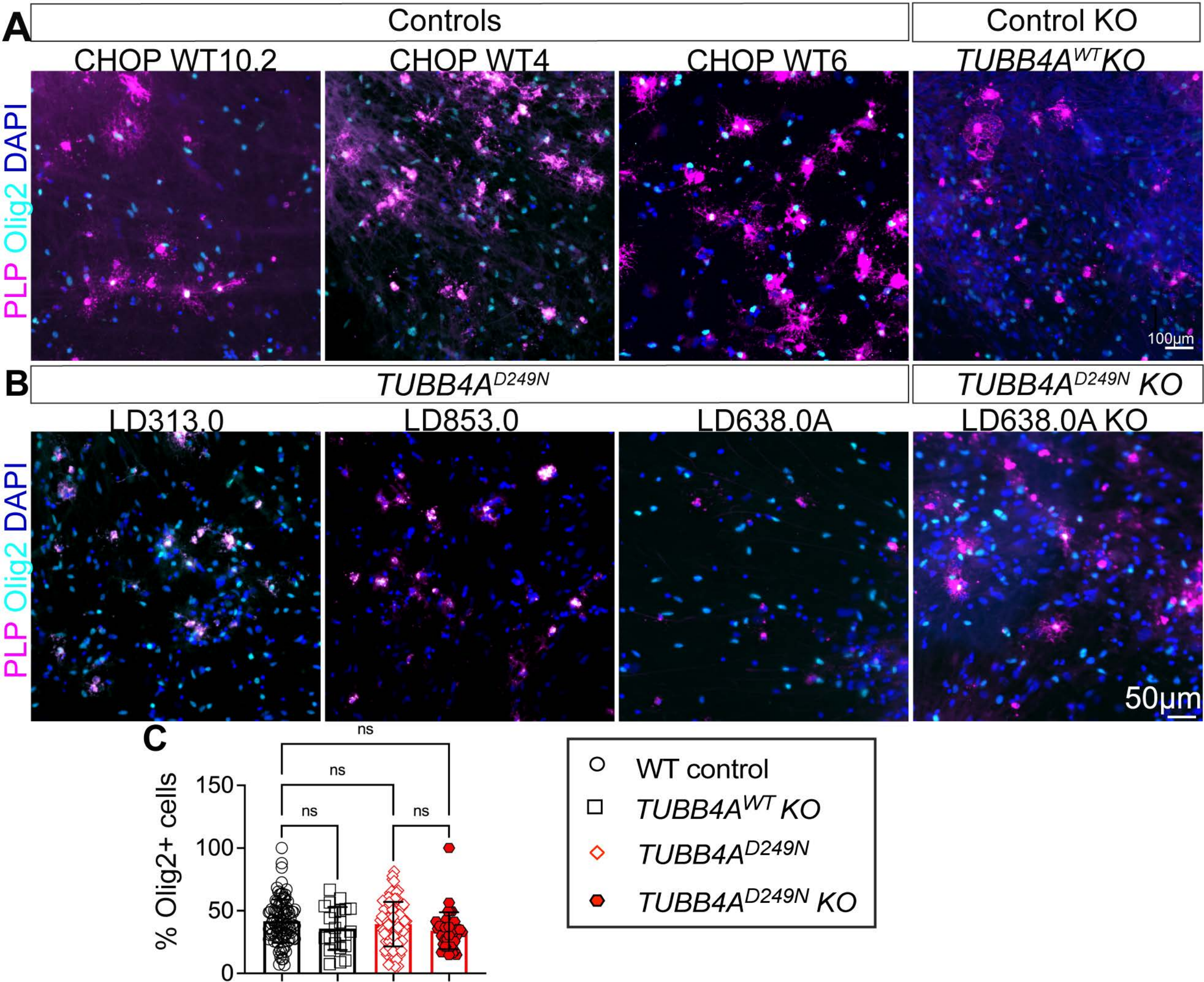

A
Controls
Control KO
CHOP WT10.2
CHOP WT4
CHOP WT6
TUBB4A^WT KO
PLP Olig2 DAPI
100µm
B
TUBB4A^D249N
TUBB4A^D249N KO
LD313.0
LD853.0
LD638.0A
LD638.0A KO
PLP Olig2 DAPI
50µm
C
% Olig2+ cells
150
100
50
0
ns
ns
ns
ns
WT control
TUBB4A^WT KO
TUBB4A^D249N
TUBB4A^D249N KO

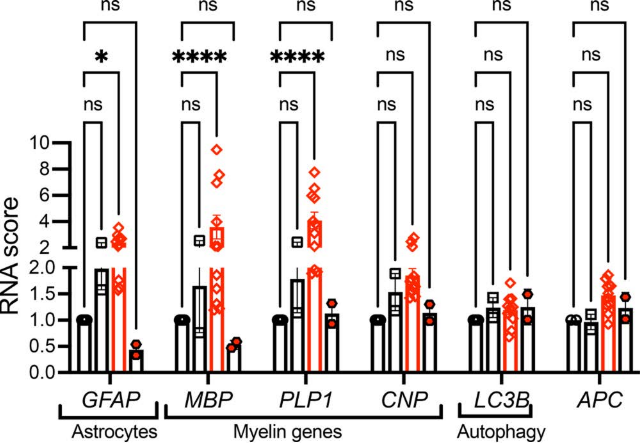
ns
ns
ns
ns
ns
ns
*
****
****
ns
ns
ns
ns
ns
ns
ns
ns
ns
RNA score
10
8
6
4
2
2.0
1.5
1.0
0.5
0.0
GFAP
MBP
PLP1
CNP
LC3B
APC
Astrocytes
Myelin genes
Autophagy

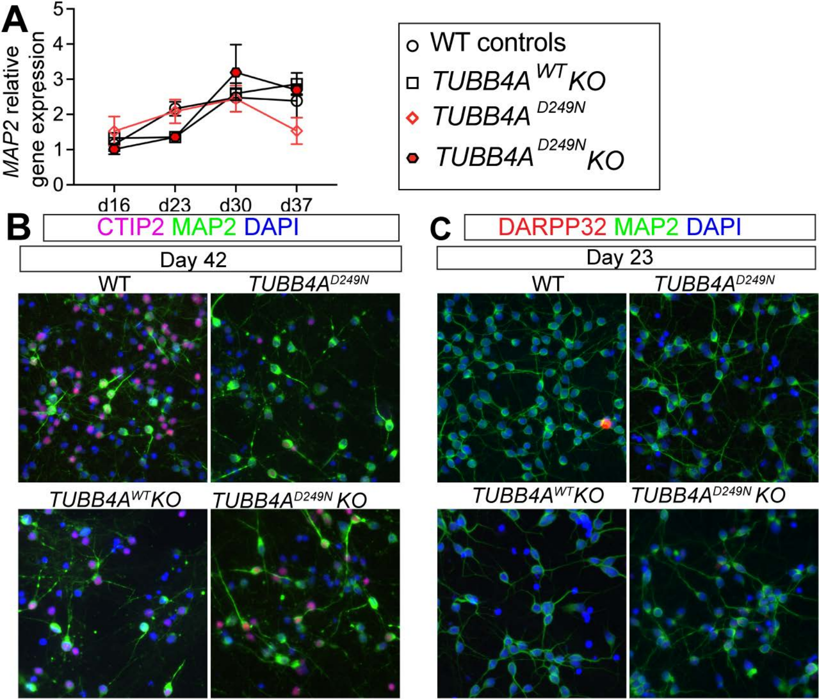
A
MAP2 relative gene expression
0
1
2
3
4
5
d16
d23
d30
d37
WT controls
TUBB4A WT KO
TUBB4A D249N
TUBB4A D249N KO
B
CTIP2 MAP2 DAPI
Day 42
WT
TUBB4A D249N
TUBB4A WT KO
TUBB4A D249N KO
C
DARPP32 MAP2 DAPI
Day 23
WT
TUBB4A D249N
TUBB4A WT KO
TUBB4A D249N KO

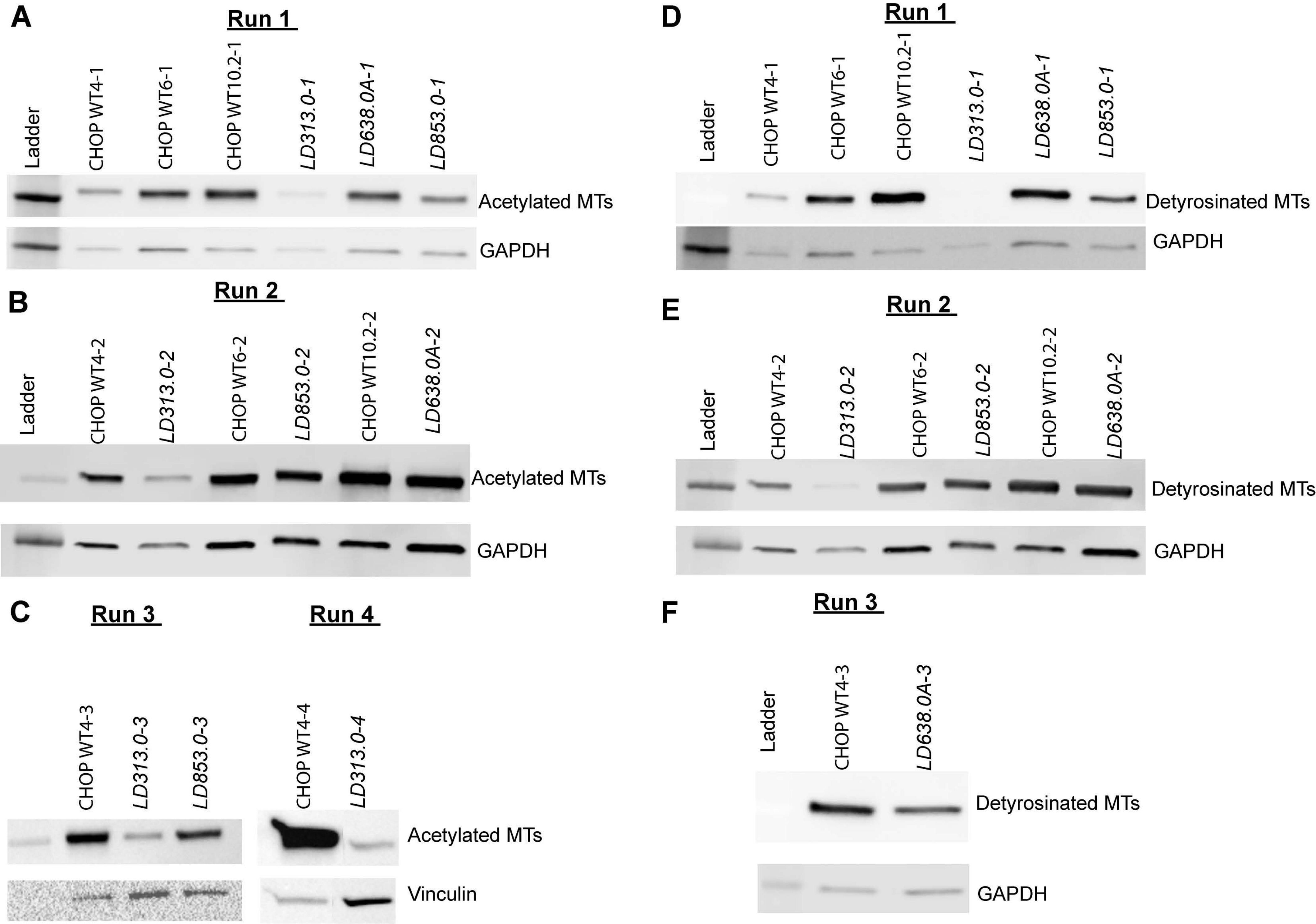

A
Run 1
Ladder
CHOP WT4-1
CHOP WT6-1
CHOP WT10.2-1
LD313.0-1
LD638.0A-1
LD853.0-1
Acetylated MTs
GAPDH
B
Run 2
Ladder
CHOP WT4-2
LD313.0-2
CHOP WT6-2
LD853.0-2
CHOP WT10.2-2
LD638.0A-2
Acetylated MTs
GAPDH
C
Run 3
Run 4
CHOP WT4-3
LD313.0-3
LD853.0-3
CHOP WT4-4
LD313.0-4
Acetylated MTs
Vinculin
D
Run 1
Ladder
CHOP WT4-1
CHOP WT6-1
CHOP WT10.2-1
LD313.0-1
LD638.0A-1
LD853.0-1
Detyrosinated MTs
GAPDH
E
Run 2
Ladder
CHOP WT4-2
LD313.0-2
CHOP WT6-2
LD853.0-2
CHOP WT10.2-2
LD638.0A-2
Detyrosinated MTs
GAPDH
F
Run 3
Ladder
CHOP WT4-3
LD638.0A-3
Detyrosinated MTs
GAPDH

Table 1

| Gender, age and source of donors of iPSC lines | | | | |
|---|---|---|---|---|
| **Genotype** | iPSC line | Gender | Age of Donor (yrs) | Source |
| **Control** | CHOP WT4 | Male | 2 | Fibroblast |
| | CHOP WT6 | Male | Unknown | Bone marrow |
| | CHOP WT10.2 | Male | 30 | PBMC |
| ***Tubb4a$^{D249N}$*** | LD_0313.0A | Male | 13.39 | PBMC |
| | LD_0638.0A | Male | 9.33 | Fibroblast |
| | LD_0853.0 | Female | 9.05 | PBMC |
| ***Tubb4a KO*** | CHOP WT6 KO | Derived from CHOP WT6 | | |
| ***Tubb4a$^{D249N}$ KO*** | LD_0638.0A KO | Derived from LD_0638.0A | | |

Table 2

| REAGENT or | SOURCE | IDENTIFIER |
|---|---|---|
| **Antibodies** | | |
| Rabbit anti-Pax6 | Thermo Fisher Scientific | Cat#42-6600; RRID: AB_2533534 |
| Mouse anti-Nestin | Abcam | Cat#ab22035; RRID: AB_446723 |
| Goat anti-Sox17 | R&D Systems | Cat#AF1924; RRID: AB_35506 |
| Mouse anti-CD184 | Stem Cell Technologies | Cat#60089; RRID: AB_2936358 |
| Goat anti-Brachyury | R&D Systems | Cat#AF2085; RRID: AB_2200235 |
| Rabbit anti-Nanog | Cell Signaling | Cat#4903; RRID: AB_10559205 |
| Goat anti-Oct-3/4 | R&D Systems | Cat#AF1759; RRID: AB_354975 |
| Goat anti-Sox2 | R&D Systems | Cat#AF2018; RRID: AB_355110 |
| Rat anti-Ctip2 | Abcam | Cat#ab18465; RRID: AB_2064130 |
| Mouse anti-Map2 | Sigma | Cat#M1406; RRID: AB_477171 |
| Rabbit anti-Darpp32 | Cell Signaling | Cat#2306; RRID: AB_823479 |
| Rabbit anti-Oilg2 | Abcam | Cat#ab109186; RRID: AB_10861310 |
| Rat anti-PLP | Obtained from Dr. Judy Grinspan | Hybridoma source |
| Rat anti-MBP | Millipore Sigma | Cat#MAB386; RRID: AB_94975 |
| Rat anti-A2B5 | Obtained from Dr. Judy Grinspan | Hybridoma source |
| Rabbit anti-detyrosinated α-tubulin | RevMAb | Cat#31-1335-00 |
| Mouse anti-acetylated tubulin | Sigma-Aldrich | Cat#T7451; RRID: AB_609894 |
| Goat anti-rabbit IgG (H+L) Alexa Fluor® 647 | Thermo Fisher Scientific | Cat# A-21245; RRID: AB_2535813 |
| Goat anti-mouse IgG (H+L) Alexa Fluor® 488 | Thermo Fisher Scientific | Cat# A-11029; RRID: AB_2534088 |

| | | |
|---|---|---|
| Goat anti-rat IgG (H+L) Alexa Fluor® 647 | Thermo Fisher Scientific | Cat# A-21247; RRID: AB_141778 |
| Donkey anti-goat IgG (H+L) Alexa Fluor® 555 | Thermo Fisher Scientific | Cat# A-21432; RRID: AB_141788 |
| Donkey anti-mouse IgG (H+L) Alexa Fluor® 647 | Thermo Fisher Scientific | Cat# A-31571; RRID: AB_162542 |
| Donkey anti-rabbit IgG (H+L) Alexa Fluor® 488 | Thermo Fisher Scientific | Cat# A-21206; RRID: AB_2535792 |
| ProLong Gold Antifade Reagent with DAPI | Cell Signaling | Cat#8961; RRID: N/A |
